# Expression of *Castanea crenata* Allene Oxide Synthase in Arabidopsis Improves the Defense to *Phytophthora cinnamomi*

**DOI:** 10.1101/2020.10.09.333781

**Authors:** Susana Serrazina, Helena Machado, Rita Costa, Paula Duque, Rui Malhó

**Affiliations:** BioISI – Biosystems & Integrative Sciences Institute, Faculdade de Ciências, Universidade de Lisboa, 1749-016 Lisboa, Portugal; INIAV—Instituto Nacional de Investigação Agrária e Veterinária, 2780-157 Oeiras, Portugal; Centro de Estudos Florestais, Instituto Superior de Agronomia, Universidade de Lisboa—Tapada da Ajuda, 1349-017 Lisboa, Portugal; Instituto Gulbenkian de Ciência (IGC), 2780-156 Oeiras, Portugal

**Author notes:** **Corresponding authors:** Susana Serrazina & Rui Malhó, BioISI – Biosystems & Integrative Sciences Institute, Faculdade de Ciências, Universidade, de Lisboa, 1749-016 Lisboa, Portugal, Address correspondence to, The author responsible for distribution of materials integral to the findings presented in this article in accordance with the Journal policy described in the Instructions for Authors (http://www.plantphysiol.org) is: Susana Serrazina.

**Keywords:** Allene oxide synthase, *Castanea crenata*, gene functional analysis, *Phytophthora cinnamomi*

## Abstract

Allene oxide synthase (AOS) is a key enzyme of the jasmonic acid (JA) signaling pathway. The *AOS* gene was previously found to be upregulated in an Asian chestnut species resistant to infection by the oomycete *Phytophthora cinnamomi (Castanea crenata),* while lower expression values were detected in the susceptible European chestnut (*Castanea sativa*). Here, we report a genetic and functional characterization of the *C. crenata* AOS (CcAOS) upon its heterologous gene expression in a susceptible ecotype of *Arabidopsis thaliana,* which contains a single *AOS* gene. It was found that Arabidopsis plants expressing CcAOS delay pathogen progression and exhibit more vigorous growth in its presence. They also show upregulation of jasmonic acid and salicylic acid-related genes. As in its native species, heterologous CcAOS localized to plastids, as revealed by confocal imaging of the CcAOS-eGFP fusion protein in transgenic Arabidopsis roots. This observation was confirmed upon transient expression in *Nicotiana benthamiana* leaf epidermal cells. To further confirm a specific role of CcAOS in the defense mechanism against the pathogen, we performed crosses between transgenic CcAOS plants and an infertile Arabidopsis *AOS* knockout mutant line. It was found that plants expressing *CcAOS* exhibit normal growth, remain infertile but are significantly more tolerant to the pathogen than wild type plants.

Together, our results indicate that CcAOS is an important player in plant defense responses against oomycete infection and that its expression in susceptible varieties may be a valuable tool to mitigate biotic stress responses.

**One-sentence summary:** Heterologous expression of the *Castanea crenata* allene oxide synthase gene in *Arabidopsis thaliana* improves the defense response to the pathogen *Phytophthora cinnamomi*.

## INTRODUCTION

The European chestnut (*Castanea sativa*) suffers significant losses in orchard production due to its most dangerous pathogen, *Phytophthora cinnamomi. P. cinnamomi* is a soil-borne hemibiotrophic oomycete that infects roots in the presence of water through motile zoospores, an infection that can also occur artificially using mycelium inocula (Moralejo et al., 2009; Redondo et al., 2015). From roots, the pathogen progresses through the vascular system, hampering water and nutrient uptake, causing host disease and eventually death (Maurel et al., 2004).

Among chestnuts, the Asian species exhibit higher resistance to *P. cinnamomi*, particularly the Japanese chestnut, *C. crenata*. We previously sequenced the root transcriptomes of *C. crenata* and *C. sativa* upon *P. cinnamomi* inoculation and found differentially expressed genes in *C. crenata* that are candidate defense genes against this oomycete (Serrazina et al., 2015). Among these, allene oxide synthase (AOS) presented a striking expression pattern, being upregulated five-fold in the resistant species while downregulated three-fold in *C. sativa* (Serrazina et al., 2015).

The defense response against soil-borne pathogens in roots begins with microorganism recognition through microbe-associated molecular patterns (MAMP) and/or plant damage-associated molecular patterns (DAMP); both can initiate a plant immune response (reviewed by De Coninck et al., 2014). Membrane-bound pattern recognition receptors of the plant recognize M/DAMPs leading to MAMP-triggered immunity (MTI) and, subsequently, cell wall fortification, production of reactive oxygen species (ROS), pathogenesis-related proteins (PR proteins) and secondary metabolites such as phytoalexins (De Coninck et al., 2014). Pathogens that can suppress MTI produce effectors that mask MAMPs, inhibit proteases and thus hamper host responses. Oomycete pathogens can also develop haustoria that are able to release effectors into the plant cell. On the other hand, plants co-evolving with pathogens can develop effector recognition (via R genes) through proteins with a nucleotide-binding (NB) and a leucine-rich repeat (LRR) domain, leading to effector-triggered immunity. This type of immunity is stronger than MTI and can give rise to the hypersensitive response, characterized by programmed cell death. After MTI, root defense responses are regulated depending on the type of threat, with the phytohormones salicylic acid (SA) and jasmonic acid (JA), among others, playing crucial roles in the primary signaling (Chen et al, 2015).

JA and derived metabolites function as signals in the response to several stimuli, including biotic or abiotic stress and wounding, as well as in developmental processes, such as pollen development and anther dehiscence, root growth or fruit ripening (Devoto and Turner, 2005). More recently, Balfagón et al. (2019) described an important role of JA in plant acclimation to intense light and heat stress.

AOS, or Cytochrome P450 74A, is predominantly localized in the plastid membrane (Froehlich et al., 2001) and takes part in the first steps of the JA signaling pathway; lipoxygenase produces 13-hydroperoxy-linoleic acid from membrane lipids, which spontaneously degrades into a keto-hydroxy fatty acid derivative that is transformed into allene oxide by AOS (Chapple, 1998). Allene oxide is rapidly cyclized by allene oxide cyclase to a more stable product, cis-(+)-12-oxo-phytodienoic acid (OPDA) (Laudert and Weiler, 1998). Our previous work pointed to the importance of the JA pathway for the chestnut defense response to *P. cinnamomi,* revealing the differential expression of both JA pathway and JA-induced genes upon *C. crenata* and *C. sativa* infection (Serrazina et al., 2015). Camisón et al. (2019) further reported that jasmonoyl-isoleucine (JA-Ile) levels increase in resistant chestnut roots following infection, while JA-Ile is practically undetectable in non-infected roots.

Despite obvious limitations in applied research, many recent plant-pathogen interaction studies have also taken advantage of *Arabidopsis thaliana,* due to the plethora of available mutants and its detailed genome annotation. Arabidopsis-Phytophthora pathosystems have been established and used to investigate the role of the JA pathway or related genes in the plant’s response to infection, namely Arabidopsis-*P. infestans* (Pajerowska-Mukhtar et al., 2008), Arabidopsis-*P. parasitica* (Attard et al., 2010) and Arabidopsis-*P. cinnamomi* (Rookes et al., 2008). Studies using the first two systems reported that Arabidopsis mutants impaired in the JA pathway exhibit enhanced pathogen susceptibility, pointing to the involvement of JA in the resistance to Phytophthora. Moreover, Attard et al. (2010) and Rookes et al. (2008) describe different responses of roots and leaves to the pathogen, suggesting that the regulation of defense genes by phytohormones is organ dependent (Chuberre et al., 2018).

In *A. thaliana*, AOS is encoded by a single gene. Given that our transcriptomic data suggested the involvement of *AOS* in the resistance of Japanese chestnut to *P. cinnamomi* (Serrazina et al. 2015), we devised a set of functional and molecular studies to test this hypothesis. We resorted to the heterologous constitutive gene expression of *C. crenata AOS* (*CcAOS*) in *A. thaliana* plants of the Landsberg erecta (L*er*-0) ecotype, known for its susceptibility to *P. cinnamomi* (Robinson and Cahill, 2003). An Arabidopsis-*P. cinnamomi* pathosystem was developed in which plants were root-inoculated under axenic conditions to analyze the response of transgenic plants expressing CcAOS. To confirm functionality of this heterologous gene expression, we performed subcellular localization analyses of CcAOS-eGFP and genetic crosses of an Arabidopsis *aos* mutant line with *CcAOS*.

We show that *CcAOS* expression in Arabidopsis is able to delay pathogen progression along the root, concomitant with an upregulation of JA- and SA-related genes in transgenic *CcAOS* plants 24 h after infection, suggesting that both signaling pathways are involved in the response to *P. cinnamomi* at early stages of host tissue invasion. CcAOS-eGFP was observed in plastids and the expression of CcAOS in *aos* loss-of-function mutants strengthens a role of the heterologous gene in plant defenses to oomycetes. Together, our results support an important role of *C. crenata* AOS in biotic stress mechanisms and open new perspectives towards the generation of new *C. sativa* cultivars.

## RESULTS

### Transgenic Arabidopsis Plants Expressing CcAOS Exhibit Slightly Accelerated Development

Arabidopsis wild type plants from the ecotype Landsberg *erecta* (L*er*-0), which are highly susceptible to the oomycete *P. cinnamomi* (Robinson and Cahill, 2003), were transformed with the *AOS* gene from *Castanea crenata* (*CcAOS*). According to our sequencing and *in silico* analysis, the *CcAOS* ORF has 1581 nt and is predicted to encode 527 amino acids, with 58.7 kDa molecular weight and 9.01 pI. After a BLASTp in NCBI, the most related sequences were found to be from *Quercus suber*, *Morus notabilis* and *Camellia sinensis*, showing 96, 80 and 76% identity, respectively (data not shown). The CcAOS amino acid sequence shares 68% identity with the *A. thaliana* AOS (At5g42650.1) (Supplemental Fig. S1). The *CcAOS* open reading frame (ORF) was cloned upstream of the eGFP sequence under the control of the constitutive CaMV35S promoter (Fig. 1A).

**Figure 1.**
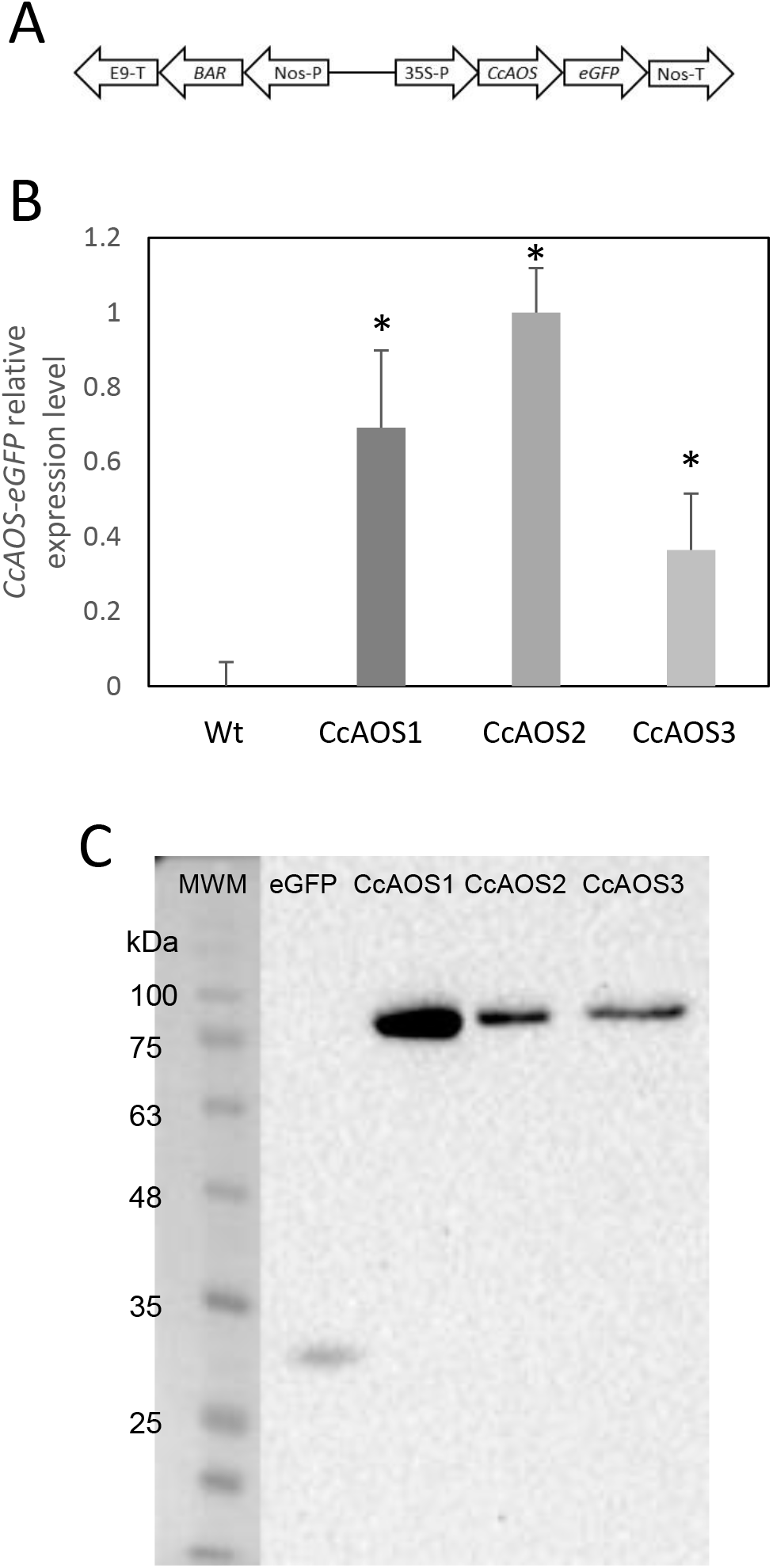
Molecular characterization of transgenic Arabidopsis plants expressing *CcAOS*. A. Schematic diagram of the *CcAOS*-eGFP construct used for transformation. 35S-P: CaMV 35S RNA promoter; *Nos-T: Nopaline synthase* terminator; *Nos-P: Nopaline synthase* promoter; *BAR: Phosphinothricin N-acetyltransferase* coding sequence; *E9-T: pea rbcS-E9* terminator. B. Relative expression levels of *CcAOS*-eGFP in two-week-old transgenic plants. Wild type L*er*-0 was used as negative control. Expression levels were normalized to the *Actin2* (At3g18780) reference gene. The highest level of expression in CcAOS2 was set to 1 and used as calibrator. Error bars represent the standard error of the mean (n=3). Asterisks indicate significant differences in expression between transformed plants (P < 0.005; Student’s *t*-test). C. CcAOS-eGFP protein expression in two-week-old transgenic plants. An eGFP monoclonal antibody was used in hybridization, and plants transformed with the empty vector were used as positive control for eGFP. Molecular weights: eGFP, 27 KDa; CcAOS-eGFP, 77 KDa. Image representative of three independent experiments.

Three independent Arabidopsis lines stably transformed with pBA-CcAOS-eGFP were isolated (Fig. 1B,C). Subsequent generations of these plants were analyzed and compared to the wild type to assess the effects of heterologous *CcAOS* expression. Transgenic *CcAOS* Arabidopsis plants were morphologically very similar to the wild type although with taller flower stalks (Fig. 2A) and larger rosettes (Supplemental Table S1). Likewise, *CcAOS* flowers displayed no visible defects (Fig. 2B), but developed earlier (Supplemental Table S1) and, upon self-fertilization, generated slightly smaller siliques with fewer seeds (Fig. 2C,D). The root system of transgenic *CcAOS* plants also appeared normal (Fig. 2E) though exhibiting longer primary root length and more lateral roots (*P*<0.05) (Supplemental Table S1).

**Figure 2.**
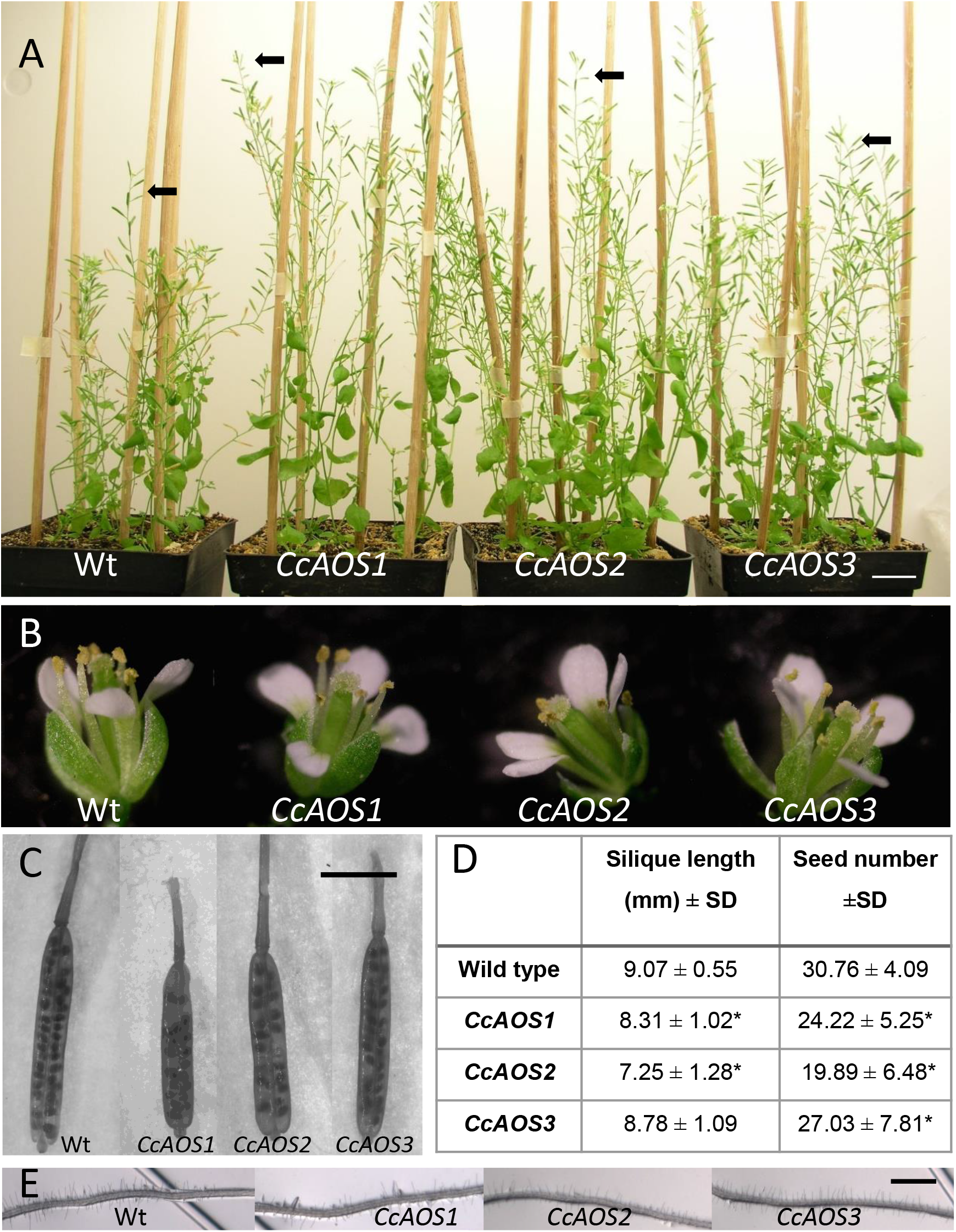
Phenotypical characterization of transgenic Arabidopsis plants expressing *CcAOS-eGFP*. A. Two-month-old wild-type (L*er*-0) and transgenic *CcAOS* plants. Arrowheads indicate the height of flower stalks. Scale bar = 2 cm. B. Detail of flowers from plants shown in A. C. Detail of siliques collected from plants shown in A. Bar = 5 mm. D. Average silique length and seed number (±SD; n=3) collected from plants shown in A. Asterisks indicate significant differences when compared to L*er*-0 plants (P < 0.05; Student’s *t*-test). E. Detail of roots from plants shown in A. Scale Bar = 100 μm.

### CcAOS Contains a Signal Peptide and Localizes to the Plastid

AOS was reported to be a plastid protein in plants (Tijet and Brash, 2002). We used an *in silico* tool to predict transit peptides in the *CcAOS* sequence, and a plastid signal peptide was found encoded in amino acids 1 to 21 (Supplemental Fig. S1). Three independent transgenic *CcAOS* Arabidopsis lines were used for subcellular localization analysis of the CcAOS-eGFP fusion protein in roots of one-week-old seedlings. Plants transformed with the pBA-eGFP binary vector alone were used as a control and compared to the three *CcAOS* lines. Observations were similar in all lines, but to reduce the possibility of artefactual localization resulting from overexpression, we focused on data stemming from line 3, which exhibited lower CcAOS levels (Fig. 1C).

As reported previously and predicted by the presence of the signal peptide, CcAOS-eGFP localization was consistent with accumulation in plastids (Fig. 3A,B) – sparse punctuated fluorescence of ~1-3 μm near the membrane of highly vacuolized cells, which were not observed in control experiments (Fig. 3C,D). This pattern is similar to observations made in *Physcomitrella patens* (Scholz et al., 2012), *Vitis vinifera* (Dumin et al., 2018) and *Camelia sinensis* (Peng et al., 2018).

**Figure 3.**
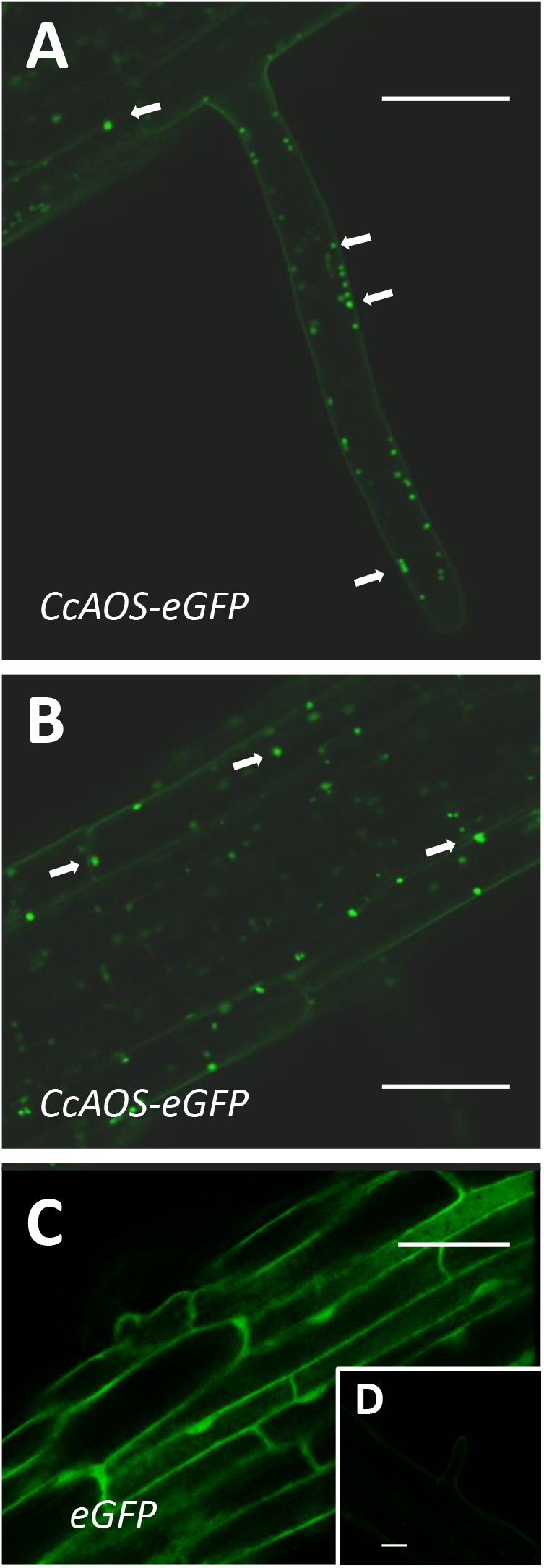
Subcellular localization of stable CcAOS-eGFP expression in roots of *A. thaliana*. Scale Bars = 30 μm. A-B. CcAOS-eGFP expression in root hairs (A) and cortical tissue of the primary root (B). Fluorescence accumulates in plastids (arrows). C. Expression of eGFP alone. D. Image of non-transformed cortical tissue.

The expression of CcAOS-eGFP in roots was significantly lower than in the aerial tissues. As imaging of the Arabidopsis small but dense leaves is recognizably difficult, we also used transiently transformed *Nicotiana benthamiana* epidermal leaf tissue for a detailed analysis of the subcellular localization and dynamics of CcAOS-eGFP fluorescence. The same pBA-CcAOS-eGFP plasmid used to transform Arabidopsis plants was infiltrated into *N. benthamiana* leaves and observations performed after four days (Fig 4). CcAOS-eGFP was found to also concentrate in punctuated structures that co-localize with chloroplasts (Fig. 4A-C) and decorate their outer regions. Interestingly, alongside with bright fluorescent spots of CcAOS-eGFP accumulation, we could also register a faint reticulate-like distribution (Fig. 4C arrows, Supplemental Fig. S2) that was not present in control experiments. This suggests that the trafficking of CcAOS to the plastids may involve and/or be mediated by the endomembrane compartment.

**Figure 4.**
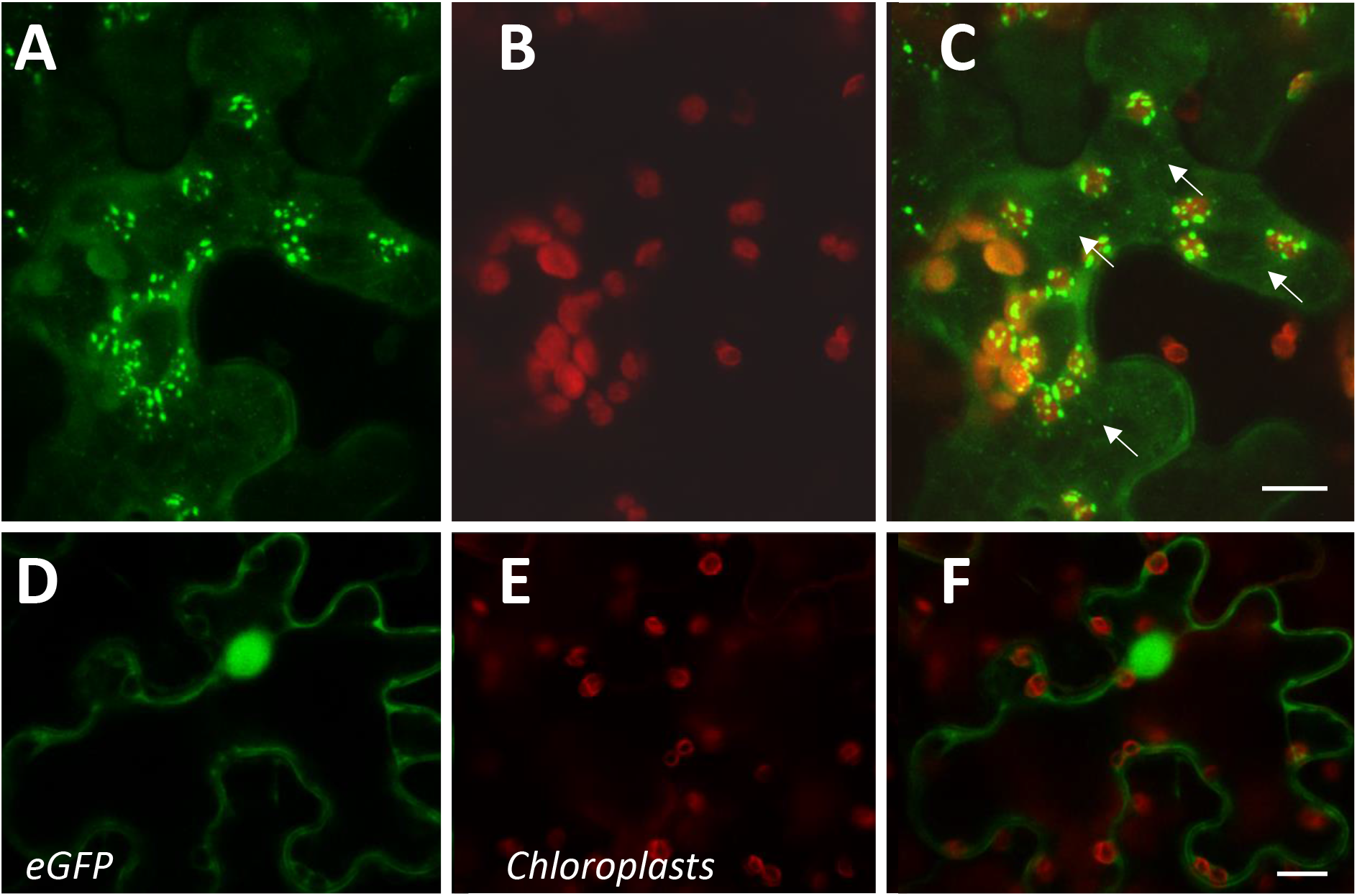
Subcellular localization of transient CcAOS-eGFP expression in leaves of *N. benthamiana*. Scale Bars = 10 μm. A-C. CcAOS-eGFP (A), chlorophyll autofluorescence (B) and merged image (C). D-F. eGFP alone (D), chlorophyll autofluorescence (E) and merged image (F).

### Expression of CcAOS in Arabidopsis Reduces Susceptibility to *P. cinnamomi* Infection and Delays Pathogen Progression

To assess if the constitutive expression of *CcAOS* affects pathogen progression along the root, we developed an axenic assay where two-week-old Arabidopsis plants growing in 0.5X MS media were inoculated at the root cap with mycelia fragments of a *P. cinnamomi* virulent strain (Fig. 5). Macroscopically, growth of mycelia along the root system was observed from 3 days after inoculation (d.a.i.) and their progression scored at 3, 6 and 9 d.a.i., after which the pathogen was able to colonize the root completely. The plants transformed with *CcAOS* showed the lowest percentage of root colonization reaching a maximum of 58% at 9 d.a.i., whereas wild type plants exhibited similar pathogen progression values at 6 d.a.i. (Fig. 5A). After colonization of the entire root, mycelia accumulated in the aerial part of the plant, with their density being notably lower in transgenic *CcAOS* plants (Fig. 5B). No necrosis was observed in roots, with leaf chlorosis being notable as soon as mycelia reached the base of the stem. No plant survival was detected two months after inoculation. In addition to physical progression along the root, we also evaluated pathogen load via quantification of a *P. cinnamomi* gene (Pyruvate, phosphate dikinase, Pdk) at days 6 and 9 after inoculation. As expected, the amount of pathogen DNA was significantly higher for wild type plants at both time points (Fig. 5C).

**Figure 5.**
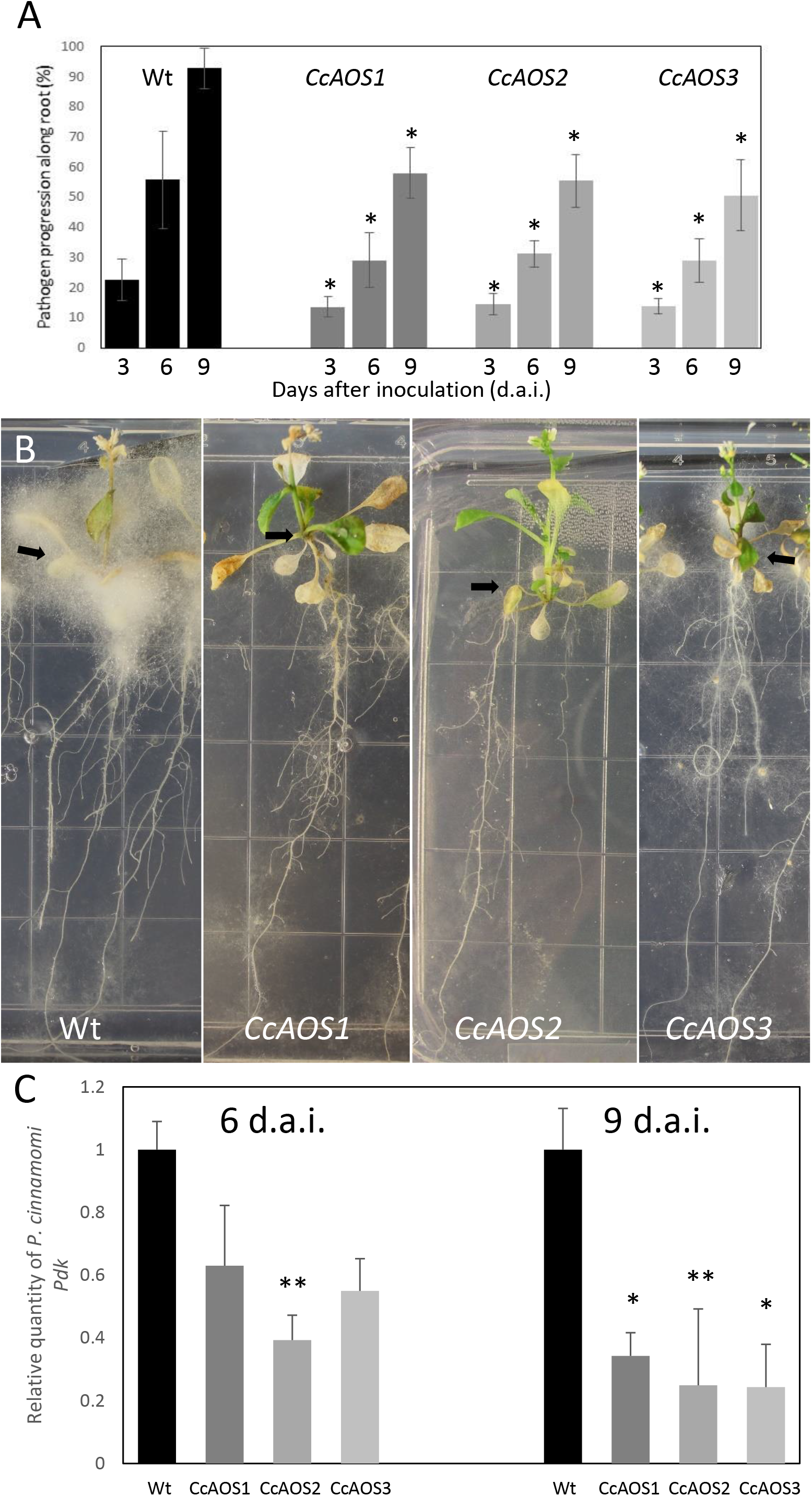
*Phytophthora cinnamomi* infection of transgenic Arabidopsis plants expressing *CcAOS*. A. Percentage of *P. cinnamomi* progression along roots of inoculated plants 3, 6 and 9 days after inoculation. Asterisks refer to significant differences from the wild-type *Ler-0* at each time point (P<0.001, Student’s *t*-test). Error bars represent the standard deviation of the mean (n=3). B. Mycelia accumulation around the plant’s aerial part (arrows) 3 weeks after inoculation. Grid squares: 1.4 cm/side. C. Relative expression of the pathogen *Pdk* gene 6 and 9 days after inoculation (d.a.i.). Values were calculated relative to L*er*-0 wild type inoculated plants and normalized to the reference genes *Mon1* and *RPF3*. Asterisks refer to significant differences from the wild type at each time-point (*P < 0.005, **P< 0.05; Student’s *t*-test). Error bars represent the standard error of the mean (n=3).

The progression of *P. cinnamomi* mycelia along roots was also cytologically followed during the first week after inoculation. Until one day after inoculation, finger-like hyphae on epidermal cells, identified as haustoria, were observed in non-transformed and *CcAOS* transformed roots (Fig. 6A, B). Haustoria are specialized hyphae capable of penetrating the host cell for nutrient uptake from the cytoplasm (Redondo et al., 2015). Two days after inoculation, hyphae reached the cortex both intercellularly and intracellularly (Fig. 6C, D), and we identified hyphal aggregations corresponding to stromata. These structures can store nutrients obtained from the host, resulting in *de novo* production of mycelium (and spores) when conditions are favorable (Willetts, 1997). At this time point, unlike with *CcAOS* plants, hyphae were frequently observed deep in the wild type root, associated with xylem vessels in vascular tissue (Fig. 6C). None of the described pathogen structures were observed in association with tissue necrosis.

**Figure 6.**
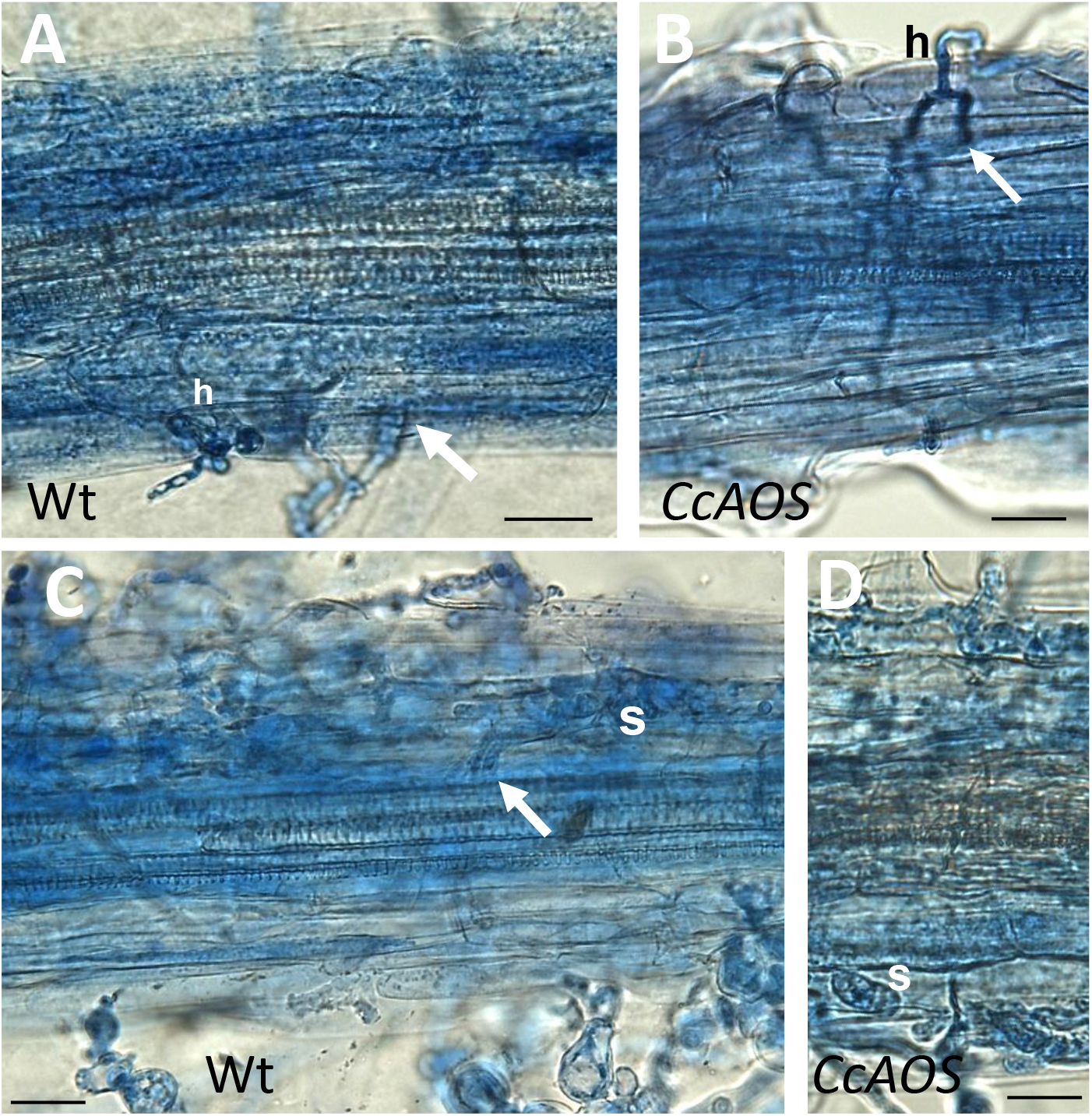
Cytological analysis of transgenic *CcAOS* Arabidopsis roots inoculated with *P. cinnamomi*. Scale Bars = 30 μm. A-B. One day after inoculation, mycelia develop haustoria (h) on epidermal cells of *Ler-0* wild type (A) and transgenic *CcAOS* roots (B). Arrows indicate hyphae penetrating the cortex intercellularly. C-D. Three days after inoculation, stromata (s) develop in cortical cells and hyphae can be observed in the stele (arrow).

These results show that the *AOS* gene from *C. crenata* delays *P. cinnamomi* progression *in planta*, when constitutively expressed in Arabidopsis. However, a direct correlation between quantity of *CcAOS* transcript or protein expressed and the effect on pathogen infection was not observed in our three plant lines. This could represent either biological or technical variability, but it is also possible that the plant’s net response reflects an effect of *CcAOS* over-expression on other metabolic parameters.

### Genes Related to Jasmonic Acid and Salicylic Acid Pathways are Upregulated in Transgenic *CcAOS* Arabidopsis Plants

The data collected supports a relevant role for *CcAOS* against *P. cinnamomi* resistance. To gain insight into the underlying defense mechanisms, we analyzed the expression of genes related to the JA and SA pathways, given their recognized involvement in biotic stresses (Armengaud at al., 2004; Clarke et al., 2000). The early defense responses of Arabidopsis against *Phytophthora* pathogens are known to differ between root and leaves (Attard et al., 2010; Robinson and Cahill, 2003; Rookes et al., 2008). We therefore evaluated differential gene regulation in root and leaf tissues. The selection of the time points — 3, 12 and 24 hours after inoculation (h.a.i.) — was based on the observation of haustoria as early as 1 day after inoculation. Haustoria are structures of cell invasion that correspond to pathogen initial biotrophic growth and, according to Attard et al. (2010), SA- and JA-signaling pathways are promptly triggered when the oomycete penetrates roots.

The expression of the endogenous *AtAOS* gene was first assessed to verify transcript fluctuations in the transgenic plant lines. Figure 7 shows a similar transcript profile in wild type and transgenic *CcAOS* plants, with a significant upregulation in leaves at 24 h.a.i. but not in roots. The similar *AtAOS* expression pattern in the wild type and in plants constitutively expressing *CcAOS*, suggests a non-deleterious effect of the heterologous protein in the morphology and development of transformed plant lines.

**Figure 7.**
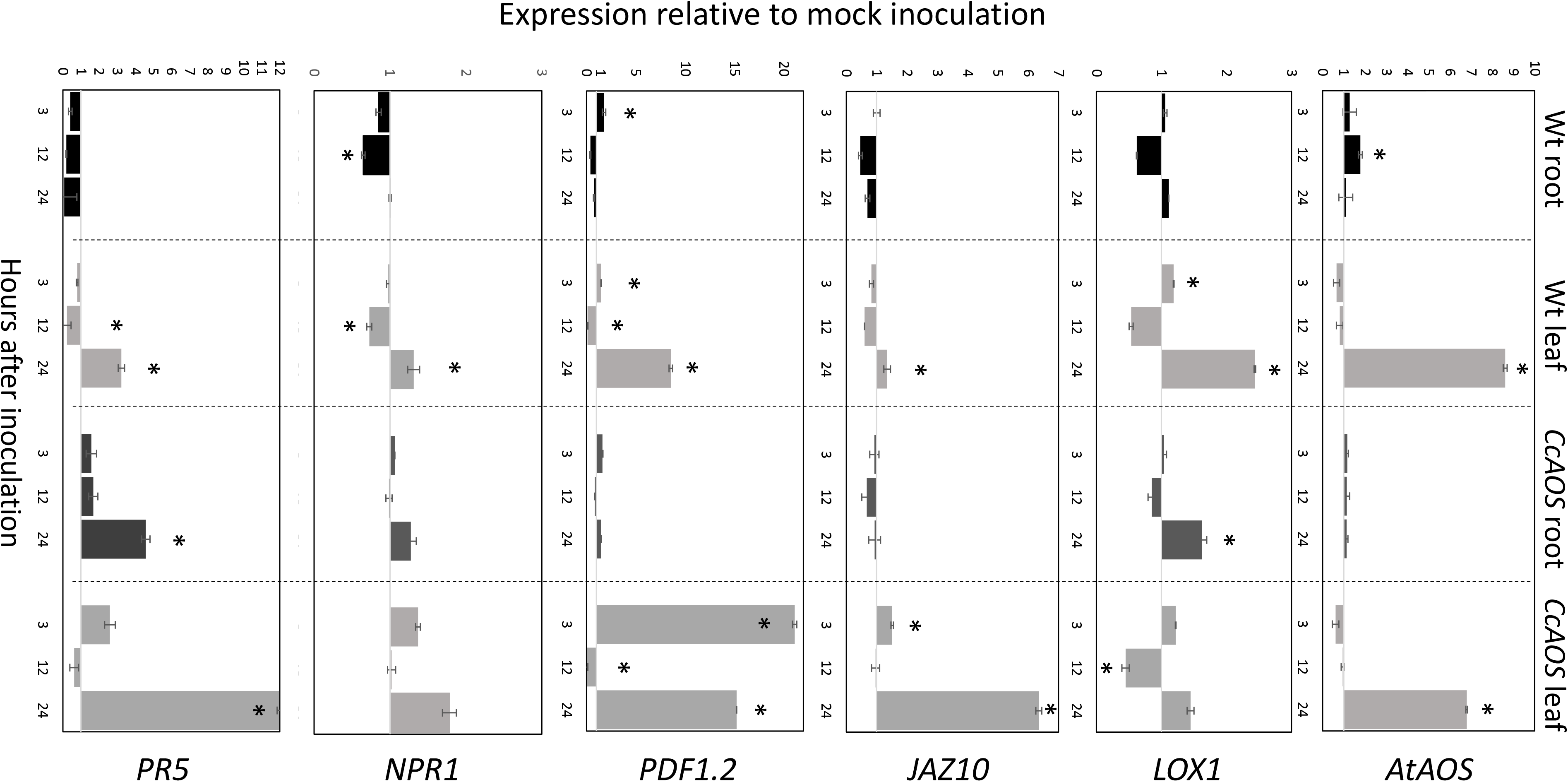
Relative expression of JA and SA pathway related genes in two-week-old transgenic *CcAOS* Arabidopsis plants at 3, 12 and 24 hours after inoculation (h.a.i.) with *P. cinnamomi*. Expression of the *LOX1, JAZ10, PDF1.2, NPR1* and *PR5* genes was calculated relative to mock-inoculated plants and normalized to the *Mon1* and *RPF3* reference genes. Error bars represent the standard error of the mean (n=3). Asterisks indicate significant differences in the expression when compared to mock-inoculation (*P*< 0.05, Student’s *t*-test).

Previously, we reported significant differential expression of two JA-marker genes, *LOX1* and *JAZ10*, in *Castanea* upon *P. cinnamomi* inoculation (Serrazina et al., 2015). Lipoxygenase-1 (LOX1) is a 9S-lipoxygenase that precedes AOS in the JA pathway and was found to play an important role in plant defense against pathogens (Vellosillo et al., 2007). Here we found that *LOX1* shows a significant upregulation only 24 h.a.i. in leaves, particularly in wild type plants (Fig. 7). By contrast, *JAZ10* exhibited a significant upregulation 24 h.a.i. in leaves of *CcAOS* plants, while in the wild type this increase was incipient. JAZ10 is a member of the JASMONATE-ZIM-DOMAIN family which reportedly negatively regulates the JA defense response, promoting growth (Guo et al., 2018). In the susceptible *C. sativa*, the *JAZ10* transcript was found to be downregulated upon *P. cinnamomi* inoculation (Serrazina et al., 2015).

A downstream marker of JA pathway, *PLANT DEFENSIN 1.2* (*PDF1.2*) was previously tested in the interaction Arabidopsis - *P. cinnamomi* (Rookes et al., 2008). Here we observed a significant upregulation of *PDF1.2* after *P. cinnamomi* inoculation in leaves of *CcAOS* plants at a very early time-point (3 h.a.i.), which was only moderately detected in wild type susceptible leaves. This further confirms the involvement of *CcAOS* and the JA pathway in the defense responses. Together, our analysis of JA-related genes suggests that the over-expression of *CcAOS* stimulates the JA pathway (downstream of *AtAOS*), activating plant defenses while alleviating growth reduction, thus allowing the plant to reach the reproduction phase.

Our previous work had also suggested that the SA pathway plays an important role in the local and systemic defense responses to *P. cinnamomi* (Serrazina et al., 2015). We thus included two SA-related genes in the analysis of transgenic *CcAOS* plants: NONEXPRESSER OF PR GENES 1 (*NPR1*) and PATHOGENESIS-RELATED GENE 5 (*PR5*). *NPR1* is referred to as a regulator of the interaction between the SA and JA pathways (Proietti et al., 2018) and has been used as a SA-marker gene in the Arabidopsis-*P. cinnamomi* interplay (Rookes et al., 2008). Here we found minor differences in *NPR1* expression between mock and pathogen-inoculated plants (less than two-fold up- or down-regulation, Fig. 7), suggesting that at the tested time-points regulation of JA signaling by SA is not achieved at the *NPR1* level. On the other hand, *PR5* encodes a thaumatin-like protein and was used by Eshraghi et al. (2011b) as a defense SA-marker gene in *P. cinnamomi*-infected Arabidopsis. Our data show that the expression of *PR5* in wild type threatened plants is downregulated in roots, exhibiting upregulation only 24h after inoculation of leaf tissues. By contrast, inoculated transgenic *CcAOS* plants exhibited *PR5* upregulation 3h after inoculation both in roots and leaf tissues. These results suggest that the constitutive expression of CcAOS in Arabidopsis promotes the expression of pathogenesis-related proteins through the crosstalk between JA and SA pathways in a *NPR1*-independent manner.

Noteworthily, despite *CcAOS* plants showing reduced susceptibility to *P. cinnamomi*, the susceptible nature of both the transgenic and wild type genotypes is reflected in the general downregulation of all analyzed genes at 12 h.a.i., which may correspond to an important stage of pathogen hijacking of JA and SA signaling through its effectors (Herlihy et al. 2019).

### The Arabidopsis and chestnut AOS fulfill distinct roles *in planta*

In Arabidopsis, loss of function of the single *AOS* gene (AT5G42650) was previously reported to cause male sterility (Park et al., 2002; von Malek et al., 2002). Here we obtained a new Arabidopsis *AOS* mutant allele, *aosGK624b02*, from the Gabi-Kat collection of T-DNA insertion mutants. Such mutant, with Col-0 background, has low susceptibility to *P. cinnamomi* (Robinson and Cahill, 2003). Consistent with the phenotype reported for the initial *aos* mutant lines (Park et al., 2002; von Malek et al., 2002), our *aosGK624b02* plants developed anthers with small filaments, incipient siliques and exhibited disturbed pollen maturation and viability (Supplemental Fig. S3). To investigate a potential role of the CcAOS protein in plant fertility, F1 plants obtained from the genetic crossing of the *aosGK624b02* mutant (pistils) and a *CcAOS* transgenic line (pollen) were let to self-pollinate and produced fertile seeds. Upon selection with two selection agents, sulfadiazine and BASTA, 28 resistant plants from F2 were obtained and grown in soil to maturity. From those, 22 displayed regular anthers that were able to generate viable pollen and to pollinate the stigma, resulting in normal siliques (Fig. 8, fertile plants). Upon genotyping, the non-disrupted *AtAOS* allele from the original transgenic pollen was detected in these plants along with the *CcAOS* transgene and the T-DNA disrupted *AtAOS* (Supplemental Fig. S4). The remaining 6 F2 plants produced small anthers with scarce pollen and abnormal siliques (Fig. 8, infertile plants). Genotyping of these plants detected the presence of *CcAOS* and the disrupted *AtAOS*, but no wild type *AtAOS* gene (Supplemental Fig. S4). Thus, heterologous expression of the *C. crenata AOS* gene was unable to restore fertility of the Arabidopsis *aos* insertion mutant, suggesting a distinct biological function from the endogenous AtAOS.

**Figure 8.**
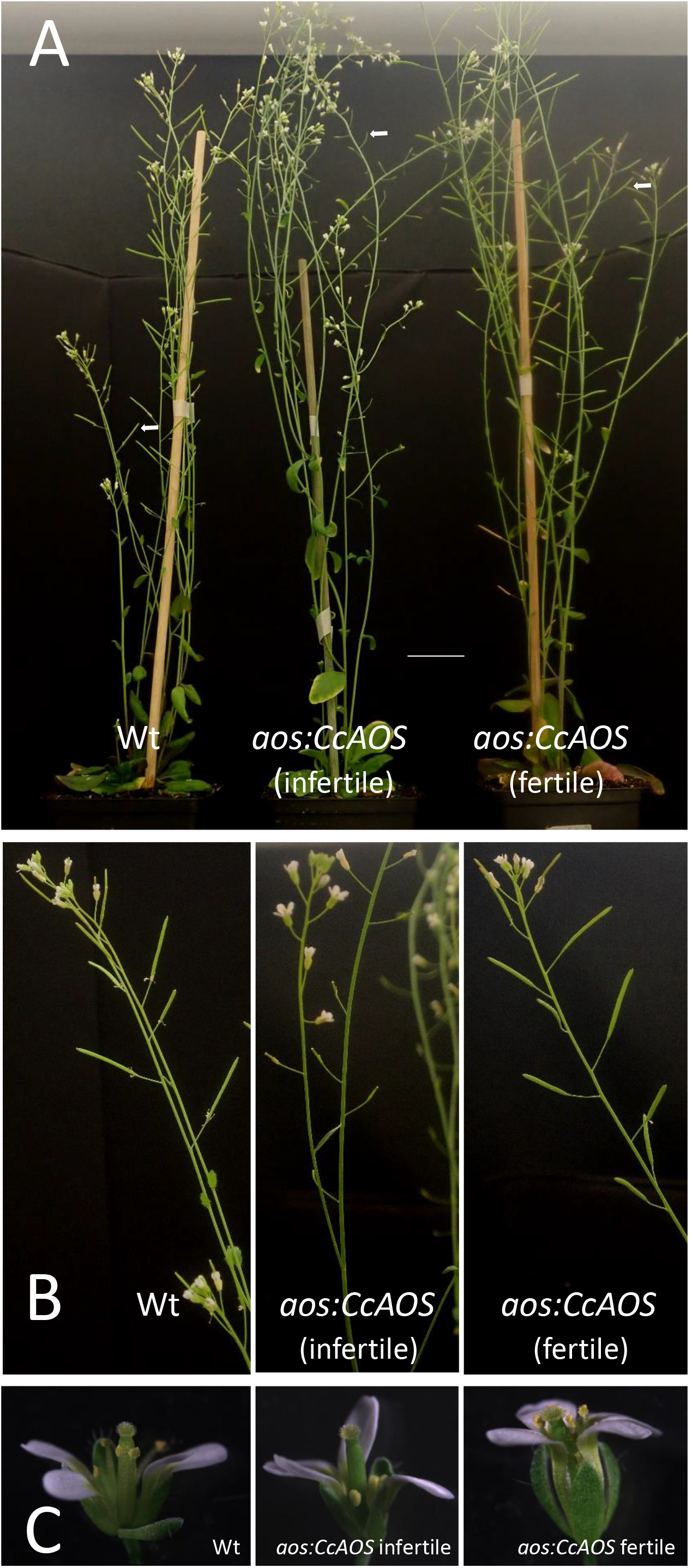
Genetic crossing of a *CcAOS* transgenic line with the Arabidopsis *aos-GK624b02* mutant. A. Two-month-old Col-0 wild-type, *aos:CcAOS* infertile (no inherited *AtAOS*) and *aos:CcAOS* fertile (inherited *AtAOS*) F2 plants. Arrows indicate siliques. Scale bar = 3cm. B. Detail of flower stalks with siliques. C. Detail of flowers.

To evaluate whether the *A. thaliana* AOS is involved in *P. cinnamomi* resistance and confirm a role for CcAOS in defense responses, we compared mycelia progression in wild type, *aosGK624b02* mutant and *aosGK624b02:CcAOS* F3 plants upon inoculation with *P. cinnamomi* (Fig. 9). After germination in selection media, two-week-old plants were inoculated and data collected during 9 days. No statistically significant differences were observed between wild type and *aosGK624b02* plants (Fig. 9A) indicating that the endogenous AtAOS does not play a role in resistance to the oomycetes. However, while a clear accumulation of mycelia was observed in roots and aerial tissues of the Col-0 wild type and the *aos-GK624b02* mutant, *aosGK624b02:CcAOS* plants exhibited a marked delay in pathogen advance on roots (Fig. 9B), clearly substantiating the notion that *C. crenata* AOS confers resistance to *P. cinnamomi* in *A. thaliana*.

**Figure 9.**
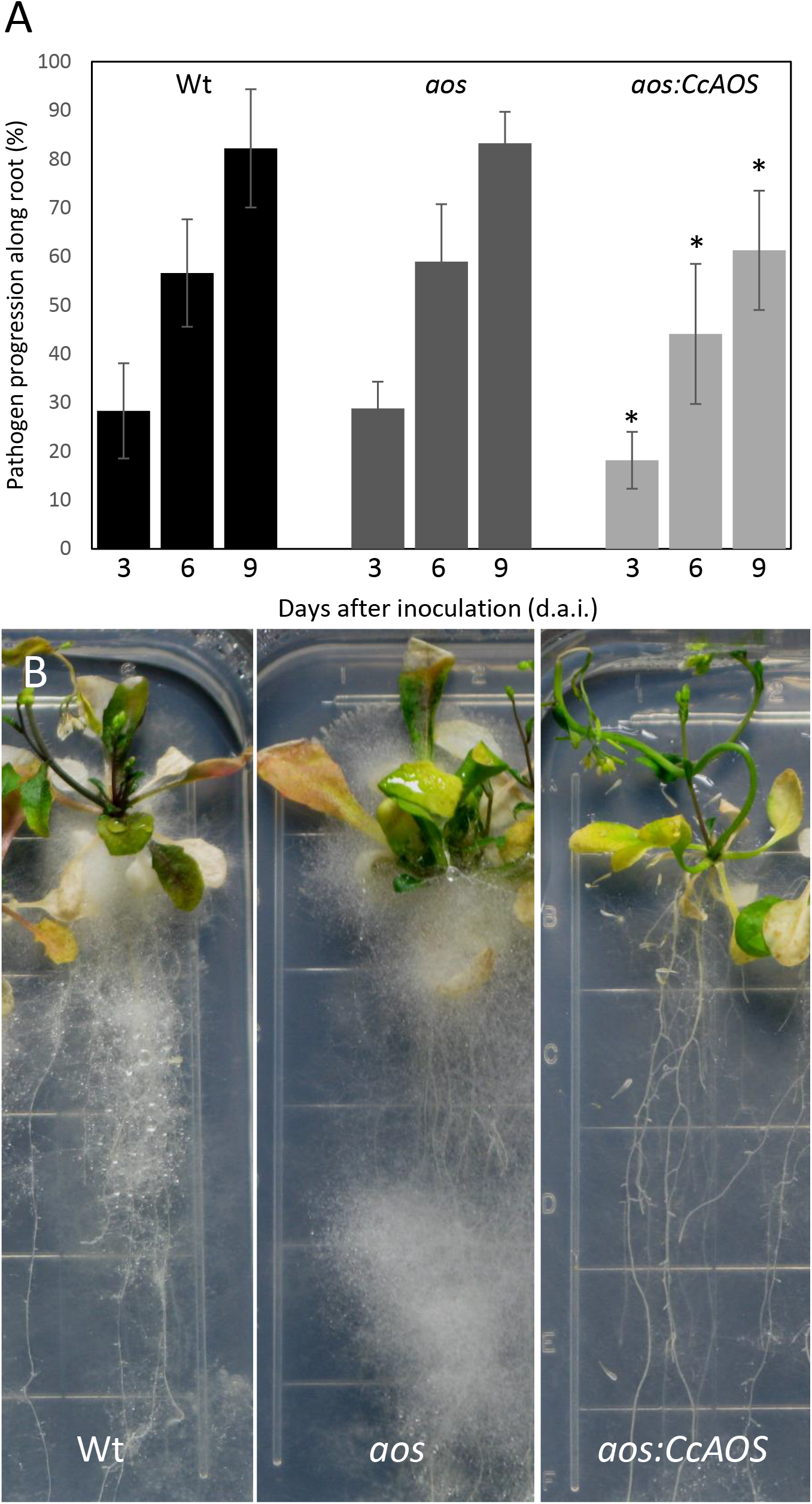
*Phytophthora cinnamomi* progression in wild type (Wt), *aos-GK624b02* mutant (*aos*) and *CcAOA*-expressing plants. A. Percentage of *P. cinnamomi* progression along roots 3, 6 and 9 days after inoculation (d.a.i.). Asterisks refers to significant differences from the Col-0 wild type at each time point (P<0.001, Student’s *t*-test). Error bars represent the standard deviation of the mean (n=3). B. Mycelia accumulation along roots and around the plant’s aerial part, 3 weeks after inoculation. Grid squares: 1.4 cm/side.

## DISCUSSION

We previously proposed an important role for allene oxide synthase (AOS) in the defense mechanisms of the Japanese chestnut, *Castanea crenata*, against the pathogenic oomycete *Phytophthora cinnamomi* (Serrazina et al., 2015). To further investigate and confirm the significance of the *C. crenata* AOS in this biotic stress response, we resorted to its heterologous expression and functional characterization using an Arabidopsis L*er*-0 ecotype-*P. cinnamomi* pathosystem. To assess correct localization of the expressed CcAOS protein, its sequence was fused with the eGFP reporter gene and fluorescence followed both in transgenic Arabidopsis plants and upon transient expression in *N. benthamiana* leaves. Furthermore, we used a loss-of-function allele for the endogenous *A. thaliana AOS* gene and its genetic crossing with the transgenic *CcAOS* Arabidopsis line to confirm a role for the chestnut AOS protein in pathogen defense and gain insight into the function of the AOS enzyme in the two plant species.

### *CcAOS* confers Oomycete Resistance to Arabidopsis Without Compromising Plant Growth and Fertility

Like the European chestnut *Castanea sativa*, Arabidopsis plants of the L*er*-0 ecotype are susceptible to *P. cinnamomi* (Robinson and Cahill, 2003). Given the amenability to transformation and functional characterization studies, we used this model flowering plant to express the *Castanea crenata* AOS gene, which we previously found to be upregulated in this species upon oomycete infection (Serrazina et al., 2015). Using the strong constitutive 35S promoter, *CcAOS* was constitutively expressed in Arabidopsis, as assessed in three independent transformed plant lines by fluorescence confocal microscopy and western blotting.

Our results show that, morphologically, transgenic *CcAOS* plants display mild differences when compared to wild type. Arabidopsis plants expressing the CcAOS transcript are distinguished by its higher growth rate, smaller siliques and lower number of seeds when compared with their WT counterparts. As previously reported by Farmer and Goosens (2019), higher basal level of JA due to an upregulation of CcAOS, resulted in plants with higher growth rate, a mechanism that could also be occurring in our heterologous lines. Interestingly, Kubigsteltig and Weiler (2003) obtained several Arabidopsis lines affected in the transcriptional control of *AOS* and, from eight lines showing constitutive *AOS* expression, all showed signs of growth inhibition, suggesting a different role for CcAOS which may account for the resistance exhibited by *C. crenata*. In two of those lines, flower development and anther size were so affected that they failed to set seed (Kubigsteltig and Weiler, 2003). In the present study, transgenic *CcAOS* plants were still able to generate viable seed set, though Kubigsteltig and Weiler (2003) did not present results on the fertilization level of the other classes of fertile mutants that would enable a comparison with *CcAOS* plants. Regardless that in all *CcAOS* plants a correlation was not evident between transcripts levels and protein expression (or with pathogen progress/quantity), in all lines isolated the pathogen progression was delayed, and the amounts of pathogen DNA were reduced when compared to non-transformed plants. Our data therefore provides compelling evidence that the *C. crenata AOS* gene positively regulates plant tolerance to oomycete infection in Arabidopsis.

The implemented Arabidopsis – *P. cinnamomi* pathosystem in axenic conditions provided us with a controlled and confined system which allows both the following of the pathogen progression and plant adaptation throughout the time. We focused on the first 5 d.a.i. for microscopic observations of root and pathogen tissues and on 12 d.a.i. for mycelia progression along the root system. After these time periods a substantial invasion of the mycelia over and inside root tissues hampered further clear observations. Despite the massive hyphal invasion, neither wild type or transgenic *CcAOS* plants showed necrotic tissues in roots. Arabidopsis ecotypes less susceptible to *P. cinnamomi*, such as Col-0, promote the formation of callose plugs in root infected areas, as well as hypersensitive response and necrotic lesions in leaves, in order to restrain the pathogen (Robinson and Cahill, 2003; Rookes et al., 2008). Here, L*er*-0 root staining with aniline blue did not reveal distinct production of callose plugs (not shown), in agreement with the susceptibility phenotype described for this ecotype (Robinson and Cahill, 2003). By contrast, the transgenic *CcAOS* plants generated in the L*er*-0 background exhibited a significant reduction in mycelia progression along roots and in *P. cinnamomi* DNA levels, demonstrating a notable contribution of the *C. crenata* AOS to plant defense.

### CcAOS Role Involves Changes in the Jasmonic and Salicylic Acid Pathways

Our results on the expression of endogenous *AOS* in inoculated L*er*-0 wild type plants agree with a susceptibility phenotype — virtually unaffected in roots and significantly upregulated in leaves 24 h after inoculation (h.a.i.). This is consistent with reports that AOS activity is highly enhanced in Arabidopsis leaves after wounding (Laudert et al. 2000). Our observation that endogenous *AOS* expression follows a similar pattern in transgenic *CcAOS* plants supports a non-deleterious effect of the heterologous protein in the morphology and development of transformed plant lines. Taken together, our data indicate that the lower susceptibility of transgenic *CcAOS* plants to the oomycetes *P. cinammoni* results specifically from the introduction (and expression) of the *CcAOS* gene and its downstream effects.

JA and SA are two of the main phytohormones implicated in plant responses against pathogens (Clarke et al., 2000; Rookes et al., 2008). Most of the studies performed so far were conducted in leaves and reported an antagonistic interaction between JA and SA signaling (Chuberre et al., 2018). However, Attard et al. (2010) reported that in roots of *Arabidopsis thaliana* both phytohormones are activated coordinately upon *Phytophthora parasitica* infection, suggesting that the early defense activation differs between roots and leaves. In the present study, we observed significant changes in the transcript levels of JA- and SA- marker genes in Arabidopsis L*er*-0 inoculated with *P. cinnamomi*. Lower expression levels were recorded in roots in agreement with Rookes et al. (2008) and Attard et al. (2010); the root was the site of pathogen inoculation and L*er*-0 plants present the most susceptible background, allowing the pathogen to efficiently suppress host immunity. The expression of *LOX1, AOS, JAZ10, PDF1.2* (JA-related) and *PR5* (SA-related) showed relevant upregulation in leaves at 24 h.a.i., revealing an amplification of the response far from the sites of inoculation (roots). Once a response to the pathogen is triggered, roots can induce a response in non-challenged organs (such as leaves), corresponding to a systemic acquired resistance (SAR, reviewed by Chuberre et al., 2018).

In roots of transgenic *CcAOS* plants, upregulation of *LOX1* expression at 24 h.a.i. was higher than in the wild type counterpart. This agrees with our predicted role of *CcAOS* in the response to biotic stress since LOX1 acts in the systemic defense against bacterial pathogens in Arabidopsis roots as shown by Vicente et al. (2012). In this report, the authors suggested that LOX1 mediates the production of oxilipins of the 9-lipoxygenase pathway which, in turn, activate SAR and regulate lateral root development (Vicente et al., 2012). In Arabidopsis roots, 9-oxilipins may activate cell wall-based responses to the fungus *Golovinomyces cichoracearum* (Marcos et al., 2015). A causal relationship between the constitutive expression of CcAOS and the higher levels of *LOX1* in roots cannot be unequivocally established; LOX1 is in the 9-lipoxygenase pathway and AOS is in the 13-lipoxygenase pathway. However, a role for oxylipins in the adaptation to adverse growth conditions and defense responses is well established (Armengaud et al., 2004) and their biosynthesis is initiated by the action of 9-LOX and 13-LOX (Vellosillo et al., 2007). Thus, a higher metabolism of common polyunsaturated fatty acids may stimulate both pathways. Moreover, LOX proteins expressed upon apple climacteric ripening were found to have dual (9/13) positioning specific lipoxygenase function (Schiller et al., 2015).

*JAZ10* upregulation in leaves of transgenic *CcAOS* plants 24 h.a.i. was also clear and in contrast with the absence of differential regulation in roots. Yan et al. (2007) suggested that JAZ10 is responsible for a repression of JA-regulated growth retardation in wounded roots of the Arabidopsis Col-0 ecotype. However, the absence of differential regulation of *JAZ10* in roots upon *P. cinnamomi* inoculation suggests an inhibitory effect of the pathogen on the defense response in its site of action. In leaves, *JAZ10* expression showed opposite regulation, in phase with the upregulation of *AOS* at the same time-point (24 h.a.i.). Leaves are directly exposed to light and, under such conditions, the JA pathway is highly inducible, with a higher expression of AOS that leads to biologically active jasmonoyl-isoleucine (JA-Ile) production (Farmer and Goossens, 2019). The availability of JA-Ile stimulates the fine modulation of the JA signaling (e.g. by JAZ10) which could promote growth and recovery from an infection scenario, giving the plant a chance of reproduction (Farmer and Goossens, 2019; Guo et al., 2018). JAZ proteins, through immunity-repression of JA pathway, promote growth to limit carbon starvation associated with strong defense responses and enable reproduction (Guo et al., 2018).

Among the analyzed genes, *PDF1.2* showed the highest up-regulation in leaves of transgenic *CcAOS* plants 3 h.a.i., suggesting that CcAOS promotes the synthesis of JA and consequently the expression of the JA-responsive gene *PDF1.2*. The regulation pattern corroborates the abundance of *PDF1.2* transcript 2.5 h.a.i. in leaves infected with *P. parasitica* (Attard et al., 2010). The noticeable upregulation of *PDF1.2* at 3 h.a.i. contrasts with the modest upregulation in wild type leaves, pointing to a relevant role of the plant defensin PDF1.2 in the defense response of *CcAOS* plants to the pathogen.

*NPR1* and *PR5* have been used as markers for SA signaling. NPR1 is associated to SAR as a negative regulator of the JA pathway (Derksen et al. 2013). In our study, the upregulation of *PR5* in wild type and transgenic *CcAOS* leaves was preceded by a weak induction of *NPR1*, suggesting that the crosstalk between JA and SA pathways is not relevant at the NPR1 level. Similarly, Eshragui et al. (2011a) reported that the application of phosphite (a systemic chemical elicitor of defense responses to *P. cinnamomi*) to L*er*-0 leaves, induced the expression of *PR5*, but not of *NPR1*. Our results corroborate such report and suggest a cooperation between the SA and JA pathways in the defense response independent of NPR1. Previously, Clarke et al. (2000) described a SA-mediated NPR1-independent resistance response that requires JA and ethylene signaling, in Arabidopsis challenged with the oomycete *Perenospora parasitica*. The most significant upregulation of *PR5* 24 h.a.i. in leaves and roots of transgenic *CcAOS* plants points to a relevant role of SA signaling in the defense response to *P. cinnamomi*. The induction of *PR5* in *CcAOS* roots, in opposition to the overall downregulation in wild type roots, indicates that the heterologous expression of CcAOS affects the regulation of the SA pathway to improve the early steps of the defense response to the pathogen.

### CcAOS Traffics to Plastids and Fulfills a Different Biological Function from AtAOS

The viability of heterologous expression of CcAOS as a tool to engineer cultivars less susceptible to pathogen attack requires confirmation that the protein is functional in pathogen defense and, inherently, that it localizes to the correct sub-cellular compartment to perform its catabolic activity. Here we addressed these two issues through expression of the *C. crenata* AOS in different *A. thaliana* backgrounds and live imaging of a CcAOS-eGFP fusion protein both in transgenic Arabidopsis plants and in *N. benthamiana* leaf epidermal cells.

Concerning the protein localization, we observed that an eGFP-CcAOS fusion protein was targeted to the expected final destination, the plastid. However, and quite interestingly, the observations performed in leaf epidermal cells of *N. benthamiana* showed trafficking through reticulate-like structures. It is tempting to speculate that the lower oomycete susceptibility induced by CcAOS results from post-translational processing of the protein through the endomembrane compartment. Endoplasmic reticulum (ER)-bodies accumulate defense proteins (Chuberre et al., 2018) and the ER is now recognizably a hub to sort proteins to plastids and mitochondria (Bellucci et al., 2018). Activation of endomembrane trafficking associated to an increase in salicylic acid levels during plant defense has also been reported (Ruano and Scheuring, 2020). Although appealing, this hypothesis clearly requires further experimental evidence.

Most significantly, when compared to the wild type plants, the expression of *CcAOS* in the *aos* mutant line resulted in a beneficial effect upon *P. cinnamomi* attack. This suggests that even in an *aos*-JA deficient background, CcAOS is sufficient to boost pathogen defense. The absence of significant differences in pathogen progression between the wild type and a newly isolated *aos* knockout, for which we confirmed previous reports of reduced fertility (Park et al., 2002; von Malek et al., 2002), indicates that the endogenous *AtAOS* does not play a significant role in the defense to *P. cinnamomi*. In support of this observation, Rookes et al. (2008) did not find differences between wild type Col-0 and the JA-biosynthesis mutants *coi1* and *jar1* upon *P. cinnamomi* root inoculation. Nonetheless, genetic crossing of transgenic CcAOS plants with the *aos* mutant also resulted in a beneficial effect upon *P. cinnamomi* attack, corroborating a role for the chestnut enzyme in the defense response to the pathogen. Noteworthily, expression of *CcAOS* in the male sterile *aos* mutant background led to plants that were morphologically similar to the wild type but still infertile. This indicates that, although functionally active in Arabidopsis, CcAOS fulfills distinct functions from the endogenous Arabidopsis AOS.

Taken together, the data presented here support the notion that CcAOS promotes resistance of the Japanese chestnut to oomycete pathogens and that its constitutive expression could be a valid tool to engineer cultivars from other species to overcome susceptibility (e.g. *C. sativa*) (Santos et al., 2017). Moreover, accordingly to Camisón et al. (2019), the expression of CcAOS seems to induce the jasmonic and salycilic acid pathways (upregulation of *LOX1, JAZ10* and *PR5*) contributing to a more efficient host response in the initial stages of *P. cinnamomi* infection without compromising growth and fertility. It further raises new questions about the evolution of plant lipid regulation and how protein function is achieved beyond catalytic activity.

It should be noted that the basis of the resistance to pathogens is multigenic and that the CcAOS gene by itself does not fully revert the susceptibility phenotype of L*er*-0. In fact, *CcAOS* plants did not show hypersensitive response or the characteristic production of callose plugs (Rookes and Cahill, 2008). Nonetheless, the signs of enhanced systemic acquired resistance from roots to leaves (suggested by the expression of *LOX1* and *PR5* in inoculated *CcAOS* plants) point to a priming of defense responses in surrounding plants through the volatile compounds from JA- and SA-pathways (Truman et al., 2007).

## MATERIALS AND METHODS

### Plant Material and Growth Conditions

The *Arabidopsis thaliana* (L.) Heynh. ecotype Landsberg erecta (L*er*-0) was used for transformation with the *C. castanea* AOS gene. For all experiments, Arabidopsis seeds were stratified in water at 4°C for 48-72h and surface sterilized for 1 min in 70% ethanol, 10 min in 30% bleach and 0.5% Tween 20, with mechanical mixing, followed by three washes with sterile distilled water. After, seeds were transferred to soil (turf and vermiculite, 3:1 mix) and periodically watered. Arabidopsis plants were grown at 22 °C and 70% relative humidity, with a 16 h: 8 h light: dark photoperiod using walk-in growth chambers (Aralab, Rio de Mouro, Portugal). Growth of Arabidopsis plants for transformation was under a 12h:12h light:dark photoperiod.

*Nicotiana benthamiana* Domin plants, used for transient transformation of leaf epidermal cells, were cultured from seed in soil (turf and vermiculite, 6:1 mix) and grown at 25 °C and 70% relative humidity, with a 16 h: 8 h light: dark photoperiod, in a walk-in chamber.

For selection of transformed seeds, stratification was followed by surface sterilization. Seeds were germinated on 9 cm diameter plates with 0.5X Murashige and Skoog medium with 1% agar, supplemented with 10 mg/L of BASTA (Glufosinate-ammonium PESTANAL®, Riedel-de Haën, Germany). After 7-14 days, all green seedlings were transferred to soil. The mutant genotype for *allene oxide synthase* (AOS, AT5G42650), acquired from GABI-KAT (GK_624b02, Kleinboelting et al., 2012) was in the Col-0 background and was cultured as above for Arabidopsis transformation, but selection was achieved with 5.0 mg/L of Sulfadiazine (Sigma-Aldrich, St. Louis, MI, USA).

For transgenic plant phenotyping, five seeds were germinated in the upper area of 100×51 mm squared plates or seedlings were transferred to soil after BASTA selection. For plant inoculation with *Phytophthora cinnamomi*, assays were performed *in vitro* and axenically as for plant phenotyping in squared plates.

The GK_624b02 *AOS* mutant line was crossed to the transgenic *CcAOS* lines, which were used as pollen donors to pistils from the mutant. Seeds from F1 and F2 were germinated in the presence of two selection agents, sulfadiazine and BASTA, and plants were transferred to soil and let to auto pollinate. F2 plants were genotyped as described below for the isolation of the transgenic *CcAOS* lines (primers for *AtAOS* and T-DNA insertion were as recommended upon https://www.gabi-kat.de/db/primerdesign.php and https://www.gabi-kat.de/faq/vector-a-primer-info.html respectively). Seeds from F3 were germinated in the same selection conditions and plants grown *in vitro* for pathogen-inoculation (see below).

Arabidopsis wild type Col-0, *aos* mutants and *aos:CcAOS* two-week-old plants inoculation with *P. cinnamomi* and mycelia progression was performed and analyzed as described below.

### Isolation and Cloning of the *Castanea crenata* Allene Oxide Synthase ORF

The nucleotide sequence of the *C. crenata* AOS transcript was obtained from the sequenced root transcriptome after *P. cinnamomi* inoculation (Serrazina et al., 2015). After a BLASTn and comparison of the sequences with highest homology, a prediction of the ORF and translation to the amino acid sequence was achieved. The CcAOS and *Arabidopsis thaliana* AOS amino acid sequence were aligned using BioEdit (https://bioedit.software.informer.com, version 7.0.5). The existence and position of the signal plastid peptide was predicted using Localizer (http://localizer.csiro.au/, Sperschneider, et al., 2017). Specific primers were designed at the 5’ and 3’ ends of the *C. crenata* AOS ORF (forward 5’-3’ ATGGCATCCACTTCTCTAGCTTTTC, reverse 5’-3’ TCAAAAGCTGGCCTTTTTGAG), which was amplified from inoculated *C. crenata* double stranded cDNA with Phusion High-Fidelity DNA Polymerase (Thermo Fisher Scientific, Waltham, MA, USA), following the manufacturers’ instructions. The expected amplification product was run in a 1% agarose gel, excised and purified with QIAquick Gel Extraction Kit (Qiagen, Hilden, Germany). The product was then cloned in pJET1.2/blunt within CloneJET PCR Cloning Kit (Thermo Fisher Scientific, Waltham, MA, EUA) and sequenced (Stabvida, Caparica, Portugal). After confirming its sequence, the *C. crenata* AOS ORF was sub-cloned in the pBA-eGFP binary vector, between XhoI and BamHI restriction sites and without the stop codon, resulting in pBA-CcAOS-eGFP. In this vector, 35S promoter drives the expression of CcAOS-eGFP and the nucleotide sequence was confirmed by Sanger sequencing.

### Plant Transformation

The pBA-CcAOS-eGFP vector was inserted in the *Agrobacterium tumefaciens* strain GV3101 and Arabidopsis L*er*-0 plants transformed following a modified flower-dip method (drop-by-drop method, Martinez-Trujillo et al., 2004). The resulting seeds were germinated in plates supplemented with BASTA for transformant screening. Leaf samples from putatively transformed 1-month-old plants were genotyped to verify the presence of *C. crenata* AOS ORF and eGFP in the genomic DNA, with KAPA3G Plant PCR Kit (Kapa Biosystems, Wilmington, MA, USA), following Section 2: Direct PCR. The primers used to amplify eGFP were: forward 5’-3’ GGGACGTCATGGTGAGCAAGG and reverse 5’-3’ CGTCCATGCCGAGAGTGATCC. Transformed plants were selected and let self-pollinate to generate F1. Plants of the F1 and F2 generation were also screened with BASTA and genotyped to confirm a stable transformation in each transformed line. Seeds, seedlings and plants derived from the F2 were used for subsequent heterologous protein localization, phenotyping and inoculation assays with *P. cinnamomi*.

For transient expression of the fusion CcAOS-eGFP protein in *Nicotiana benthamiana*, 5-6-week-old plants were used and leaf infiltration with Agrobacterium strain GV3101 (Sparkes et al., 2006) harboring pBA-CcAOS-eGFP or the empty pBA-eGFP vector performed.

### Subcellular Localization of the *Castanea crenata* AOS protein in Arabidopsis

Roots of one-week-old Arabidopsis lines transformed with CcAOS-eGFP or with the corresponding empty vector were observed in a Leica SP-E confocal microscope (Leica Microsistemas, Carnaxide, Portugal) with the settings described below and the 488nm laser line.

In transiently transformed *N. benthamiana*, sections of leaves from the infiltrated areas were observed 2, 3 and 4 days after infiltration. Imaging was achieved in a Leica SP8 confocal microscope (Leica Microsistemas, Carnaxide, Portugal). Optical sections (ca 2 μm thick) were acquired using a x63 ACS APO water objective (NA=1.15), <10% laser intensity (488 nm and 552 nm laser lines) and operating in the mode 1072 x 1072, 600 Hz (c. 0.3/s per frame). Image or Z-stack acquisition was linear, ensuring no signal bleed through.

Images were processed using the Image J software (https://imagej.nih.gov/ij/). All observations were repeated twice with at least 3 individual plants per genotype.

### Phenotypical Analysis of Transgenic Arabidopsis Lines

Plants were grown as described above, with growth and root parameters being measured on images of 1-month (potted plants) and 13-day old plants (*in vitro* plants), respectively, using the Image J software. Five plants per genotype were considered.

Flowering time, silique length and silique number were scored in five plants per genotype, with three flowers/siliques being collected per plant.

Results represent means of three independent assays, and Student’s *t*-test was used for statistical analysis.

### Analysis of *Castanea crenata AOS* Expression Levels in Transgenic Arabidopsis Lines

Total RNA was isolated from plants of wild type or CcAOS transformed lines from six plants (of each condition) using the RNeasy Plant Mini Kit (Qiagen, Hilden, Germany), followed by treatment with Turbo DNase kit (Thermo Fisher Scientific, Waltham, MA, USA). Three biological replicates per condition were prepared. Total RNA (3.6 μg) was used as template for reverse transcription with RevertAid H Minus Reverse Transcriptase (Thermo Fisher Scientific, Waltham, MA, USA) and primed with an oligo(dT) primer. Specific primers for *CcAOS* were designed with PrimerSelect 5.03 (DNASTAR Inc., Madison, WI, USA) (forward 5’-3’ CACGCGTCGATTTATTGTCC and reverse 5’-3’ TTTGGTGGGTTCGGCTTGTT).

Each cDNA was diluted 1:40 and 4 μL (18 ng) used per reaction, in a 25 μL final volume using Maxima SYBR Green qPCR Master Mix kit (Thermo Fisher Scientific, Waltham, MA, USA). A final concentration of 0.2 μM of each primer was used in a StepOne Real-Time PCR system (Applied Biosystems, Foster City, California, USA). Quantitative PCR (qPCR) reactions started with a denaturation step at 95°C for 10 min followed by 40 cycles of denaturation at 95°C for 15 s and annealing temperature (60 °C) for 30 s. Each set of reactions included a no template control and two technical replicates. Dissociation curves were used to analyze nonspecific PCR products.

To normalize expression data, *ACTIN2* was used. Oligos were forward 5’-3’ GGTATTGTGCTGGATTCTGG and reverse 5’-3’ CGCTCTGCTGTTGTGGTGA. Annealing temperature was 60 °C. 18srRNA was also used for normalization with identical results to *ACTIN2*. Oligos were forward 5’-3’ AGTCGGGGGCATTCGTATTT and reverse 5’-3’ ATCCCTGGTCGGCATCGTTT.

Gene expression was calculated using the ΔΔCT method (Schmittgen & Livak, 2008). The highest level of expression in transformed line CcAOS2 was used as calibrator (set to 1) to which all the other samples were compared. Student’s *t*-test was used for statistical analysis.

### Analysis of *Castanea crenata AOS* Protein Levels in Transgenic Arabidopsis Lines

Two-week-old plants transformed with CcAOS or the empty vector were first checked for the presence of AOS-eGFP or eGFP expression with a Olympus BX51 fluorescence microscope (Labocontrole, Lisbon, Portugal) equipped with a 470-490/DM505/LP515 filter, and a TIS 2MP DFK23U274 RGB camera (Infaimon, Aveiro, Lisbon, Portugal). A maximum of 100 mg of plants from each genotype was stored in triplicate at −80 °C. Tissue was grinded with liquid nitrogen before the addition of 300 μL of lysis buffer. RIPA buffer (radioimmunoprecipitation assay buffer; https://www.abcam.com/protocols/sample-preparation-for-western-blot) was used to obtain a protein extract with plastid proteins. Protease Inhibitor Cocktail (Sigma-Aldrich, St. Louis, MI, USA) was added freshly to the lysis buffer as recommended.

The total protein in each extract was quantified using the Bio-Rad Protein Assay (Bio-Rad, Hercules, CA, USA), and 24 μg run in a 10% SDS-PAGE. A monoclonal antibody for eGFP (Roche, Basel, Switzerland) was used for Western blotting analysis at 1:10000 dilution, followed by Peroxidase Affinipure Goat Anti-Mouse IgG (Jackson Immunoresearch, Ely, UK) at 1:10000, before detection with NZY ECL Supreme HRP substrate (NZYtech, Lisboa, Portugal).

### *P. cinnamomi* Inoculation Assays

Liquid cultures of *P. cinnamomi* (isolate IMI 340340 from the University of Trás-os-Montes and Alto Douro) were prepared in 50 mL of 1% PDB before incubation in a rotary shaker at 24 °C and 100 rpm for 48 h. Mycelia were then resuspended in 5 mL of 1% PDB and blended at high speed for 1 min. The concentration of mycelial fragments in the stock suspension was quantified with the aid of a hemocytometer and the mycelial suspension adjusted with 1% PDB to reach final concentration of approximately 1.0×10^4^ fragments/mL.

Two-week-old wild-type L*er*-0 and transgenic *CcAOS* lines, wild-type Col-0, *aos* mutants and F3 *aos:CcAOS* Arabidopsis plants, growing axenically *in vitro*, were inoculated at the root cap with 10 μL of mycelia fragment suspension. In mock inoculations, potato dextrose broth 1% was added. Plates were covered with a black cloth overnight to promote infection. The following day, the cloth was removed, the plates were shaded in the root area and placed vertically.

Photographs were taken at 3, 6 and 9 days after inoculation and measurements of mycelia progression along root performed using the Image J software (https://imagej.nih.gov/ij/). Mycelia progression was also observed microscopically at 1, 2, 3, 4, 5 and 6 days after inoculation; oomycete tissues in Arabidopsis roots were dyed with trypan blue 0,05% in a solution of lactoglycerol [lactic acid, glycerol, and water (1:1:1)] for 5 min and rinsed in lactoglycerol before observation in an Olympus BX51 microscope coupled with a TIS 2MP DFK23U274 RGB camera.

At least 10 plants per genotype were analyzed in 3 independent inoculation assays.

### Quantification of *in Planta P. cinnamomi* Growth

Genomic DNA was isolated from all Arabidopsis genotypes at 6 and 9 days after inoculation using the DNeasy Plant Mini Kit (Qiagen, Hilden, Germany).

Quantification of *P. cinnamomi* growth *in planta* was achieved through the quantification of the *Pyruvate, phosphate dikinase* gene (*Pdk*, GenBank assession FJ493007.1) by qPCR, based on Eshraghi et al. (2011a). Primers for *Pdk* were forward 5’-3’ GACGAGAGCGAGACAAGAA and reverse 5’-3’ CAAACGCACAAACGCACAC, and the melting temperature was 58 °C. The amount of genomic DNA used per reaction was 1.84 ng and the reaction mix is described above using 3 biological replicates (5 plants per replicate) and 2 technical replicates. *Monensin sensitivity 1 (Mon1*, At2g28390.1) of the SAND family protein served as a reference gene (Schlaeppi et al., 2010), using primers forward 5’-3’ GTGGCGGCGATGATAATGAT and reverse 5’-3’ CTAGTTCCCGCCACACCTT. *RNA Processing Factor 3 (RPF3* At1g62930.1) was also used for normalization (Czechowski et al., 2005) yielding identical results to *Mon1*. Oligos were forward 5’-3’ GAGTTGCGGGTTTGTTGGAG and reverse 5’-3’ CAAGACAGCATTTCCAGATAGCAT.

*P. cinnamomi* biomass in inoculated plants was calibrated with the level of inoculated wild type L*er*-0 plants. The experiment was repeated in three independent assays, and Student’s *t*-test was used for statistical analysis.

### Expression Analysis of Genes Related to the Jasmonic and Salicylic Acid Pathways

Total RNA was isolated from root and aerial tissue of wild type L*er*-0 and transgenic *CcAOS* lines at 3, 12 and 24 hours after inoculation, using the RNeasy Plant Mini Kit (Qiagen, Hilden, Germany), followed by treatment with Turbo DNase kit (Thermo Fisher Scientific, Waltham, MA, USA). Total RNA, cDNA, reaction reagents and cycling were as described above, and 5.9 ng of cDNA were used per reaction. Primer sequences for each gene are listed in Supplemental Table S2.

Three biological replicates (5 plants per replicate) and 2 technical replicates were analyzed. To normalize expression data, *Mon1* and *RPF3* were used, and gene expression levels were calculated as described above and calibrated with the respective mock-inoculated sample at each given time-point after inoculation. Results are from three independent assays, and Student’s *t*-test was used for statistical analysis.

## ACKNOWLEDGMENTS

The authors are grateful to N.-H-Chua for the pBA-eGFP binary vector, V. Nunes (IGC) for supervising and assistance with Arabidopsis transformation, E. Novo-Uzal (IGC) for supervising and assistance with western blotting, A.B. da Silva (FCUL) for advising on Arabidopsis phenotyping, T. Braga (FCUL) for assistance with *N. benthamiana* imaging and A. M. Fortes (FCUL) for critical reviewing of the manuscript. Imaging was performed at the Faculty of Sciences of the University of Lisbon’s Microscopy Facility, a node of the Portuguese Platform for BioImaging (PPBI-POCI-01-0145-FEDER-022122).

